# Acclimation kinetics of the holoparasitic weed *Phelipanche ramosa* (Orobanchaceae) during excessive light and heat conditions

**DOI:** 10.1101/2023.11.30.569309

**Authors:** Olivier Dayou, Guillaume Brun, Charline Gennat, Susann Wicke

## Abstract

Holoparasitic plants, such as broomrape, have abandoned a photosynthesis, relying entirely on the resources of host plants. This departure from an autotrophic lifestyle necessitates significant genetic and metabolic adaptations, offering a unique model system to elucidate responses independent of canonical plastid functions in green plants. In this study, we examined the acclimation kinetics of the holoparasitic weed *Phelipanche ramosa* (broomrape) under unfavorable temperature and excessive light conditions through a comprehensive time-course analysis of RNA sequence data and physiological monitoring. Our work unveils that suboptimal abiotic conditions induce transcriptional changes in the parasitic plant, involving coordinated expression of nuclear and plastid-encoded genes. Notably, magnesium transporters, critical for heat-induced chlorophyll conversion, were enriched among heat-repressed genes. Additionally, multiple copies of chloroplast-targeted DnaJ proteins, responsible for maintaining CO_2_ assimilation capacity in non-parasitic plants, were identified. Comparative expression analysis with the parasite’s host plants, tomato and *Arabidopsis*, revealed distinct patterns for certain plastid genes in *Phelipanche*. Furthermore, an elevation in reactive oxygen species (ROS) in the parasite coincided with the upregulation of numerous heat shock protein (HSP) genes, including HSP21, which associates with thylakoid membranes in photosynthetic plants; noteworthily, thylakoids are absent from *Phelipanche*’s plastids. Collectively, our findings suggest that plastids of the nonphotosynthetic model plant retains their ancestral role as environmental sensors. This research opens new avenues for functional-genetic research into the nuanced roles of plastids in the lifecycles of parasitic plants.

## Introduction

Plants confront a diverse array of environmental challenges, encompassing intense or fluctuating light and varying temperatures. A pivotal player in the plant stress response and acclimation is the plastid (Crosatti et al. 2013; Kleine et al. 2021). These organelles, integral to photosynthesis and many other cell functions of plants, sense concurrent environmental conditions and disturbances. In response, plants initiate specific anatomical or biochemical modifications, such as relocating plastids away from heat, light, and cold sources. Plastids, through organelle-based synthesis of phytohormones and secondary metabolites, as well as retrograde and ROS signaling, contribute significantly to pathogen defense in both pathogen-associated molecular pattern (PAMP)-triggered immunity and effector-triggered immunity (Jones and Dangl 2006; Serrano et al. 2016).

In adverse abiotic conditions like heat, plants accumulate chaperones such as heat shock proteins (HSPs) and proteins involved in signal transduction pathways. This response triggers antioxidant biosynthesis, reactive oxygen species (ROS) control, and the regulation of gene expression (Beltrán et al. 2018). Beyond a plant species-specific environmental tolerance, typically between 20-30°C, excessive heat leads to a rapid decline in photosynthetic efficiency. Consequently, photorespiration increases, photosystems shut down, thylakoid membranes become leaky, ATP flow disrupts, CO_2_ fixation by RuBisCO slows, and ROS formation intensifies (Yamori et al. 2011; Dopp et al. 2021). In the face of adverse conditions, detoxification of ROS becomes a primary process, particularly when coupled with biotic stressors. The interaction between responses to biotic and abiotic conditions can influence plant–pathogen interactions, potentially leading to a trade-off (Pandey et al. 2015; Choudhury et al. 2017).

Certain plants unable to photosynthesize exhibit aberrant plastid biology. Despite appearing as outliers in the plant kingdom, non-photosynthetic parasitic plants have significantly contributed to advancing our understanding of plant molecular processes and molecular evolution. Evolving a range of unique adaptations, parasitic plants serve as crucial model systems for studying plastid gene and genome evolution under relaxed selectional constraints (Wicke et al. 2013, 2016; Wicke and Naumann 2018). They provide insights into *de novo* gene evolution, sub- or neofunctionalization (Ichihashi et al. 2015; Yang et al. 2015; Yoshida et al. 2019), and ecological and molecular-evolutionary feedback loops (Chen et al. 2020; Lyko and Wicke 2021). These plants also display molecular plasticity, exhibiting dramatic alterations in gene expression and metabolic pathways in response to environmental changes. The ability to parasitize other plants has allowed their plastids to evolve under relaxed selectional constraints, eliminating the need to protect the photosynthetic apparatus. Noteworthily, whole-genome sequencing of various independently evolved parasitic plant species has revealed enriched gene losses in defense and stress response in various Orobanchaceae species and *Cuscuta* spp. (Sun et al. 2018; Vogel et al. 2018; Yoshida et al. 2019; Cui et al. 2020; Xu et al. 2022).

Here, we aimed to provide a first look into a holoparasitic plant’s strategies to acclimate to adverse conditions. Using the broomrape species *Phelipanche ramosa* infecting tomato or *Arabidopsis*, we first assessed the physiological response of the holoparasite when it suddenly experiences high temperatures or excessive light. A time-structured expression analysis was then employed to explore *Phelipanche’s* transcriptional changes of an exposure to heat and light stress. Our results implicate that despite its non-photosynthetic lifestyle and full dependence on the host for survival, the holoparasite retains the ability to deal with adverse conditions through plastid-mediated responses. We gathered evidence of acclimation responses that are independent from canonical key functions of the chloroplast in fully autotrophic plants. Therefore, this study’s focus on holoparasite biology is a promising start to identifying novel, very basal modulations in molecular acclimation processes in plants.

## Material & Methods

### Plant material

All host-parasite experiments were carried out using the well-established tomato-broomrape pathosystem (Brun et al. 2023), co-cultivated in semi-*in vitro* rhizotrons. Seeds of *Solanum lycopersicum* ‘Zuckertraube’ (Demeter quality) were purchased as precision germplasm (accession: G427) from Bingenheimer Saatgut AG (Echzell, Germany). Tomato seeds were germinated for seven days on Petri plates on moist filter paper, and primary-leaf developing tomato seedlings were then transferred onto GF/A Whatmann filter paper in vermiculite-packed rhizotrons. Tomato plants were then grown under greenhouse conditions (23°C, 16h/8h photoperiod, 100 µmol m^-2^ s^-1^) for 21 days, watered regularly with 0.5X Hoagland solution.

*Phelipanche ramosa* seeds were kindly provided by the Delavault lab (University of Nantes, France) and were harvested in a winter oilseed rape field in Sainte-Maxire (Deux-Sèvres, France) in 2017. Seeds were surface-sterilized in commercial bleach diluted to 4% active chlorine for five minutes and pre-washed three times in autoclaved deionized water for 30 seconds each. The seeds were then rinsed in autoclaved deionized water for another three times of five minutes each. Sterilized seeds were resuspended in an incubation medium (10 mg/ml), composed of 0.1 % (v/v) Plant Preservative Medium (PPM, Plant Cell Technology, USA) and 1 mM HEPES buffer (pH 7.5). The resuspended seeds were transferred in a culture flask wrapped with aluminum foil, and incubated at 21 °C for seven days.

Seeds of *Arabidopsis* Col-0 accession and *cp31a-1* T-DNA insertion mutant (Tillich et al. 2009) were a kind gift from the Schmitz-Linneweber lab, Humboldt-Universität zu Berlin, Germany. Seeds were sown on a mixture of soil and vermiculite (2:1) and incubated for five days at 4°C in the dark to synchronize germination. Seedlings were then grown at 21°C in a 16-h-dark/8-h-light cycle (100 µmol m^-2^ s^-1^). Four-week-old plants were transferred to rhizotrons.

### Host infection

To synchronize germination, we sprinkled the roots of 21 days old tomato plants with *rac*-GR24 (10^−6^ M) with Pasteur pipettes prior to broomrape infection. Pre-conditioned *P. ramosa* seeds were then aligned on to the tomato roots using a soft paintbrush. Rhizotrons were covered with aluminum foil, and tomato-*Phelipanche* plants were grown undisturbedly under greenhouse conditions as described above with weekly watering using Hoagland 0.5X. All experiments were conducted five weeks after infection, at which time parasite tubercles reached an advanced spider stage.

### Heat and high irradiance treatments

Parasite-host co-cultures were subjected to either 42°C or 1400 µmol m^-2^ s^-1^ light intensity for 4.5h then returned to control conditions (23°C, 100 µmol m^-2^ s^-1^). Rhizotron lids were removed and rhizotrons were covered with one layer of saran foil to prevent water evaporation. Heat-stressed rhizotrons were additionally covered with one layer of aluminum to avoid direct exposure to light. Parasite tubercles were carefully detached from host roots and immediately snap frozen in liquid nitrogen to quench metabolic activities. Tubercles grown in control conditions were used as reference.

### Photosynthesis measurement

Photosynthesis efficiency - also referred to as Y(II) - of tomato leaves was assessed using a Maxi-Imaging Pulse-Amplitude Modulation (PAM) fluorometer (Heinz Walz, Germany). Three plants per species were randomly selected for PAM measurements per condition per timepoint. Plants were first dark-adapted for 15 minutes to deoxidize photosynthetic reaction centers and relax energy-dependent quenching processes. For all experiments, we used a measuring light intensity of 1 Hz and a saturation pulse intensity of 7 for 200 ms. Capturing plant responses was performed in live video mode. In fluorescence mode, we selected up to three leaflets per tomato and manually delimited areas of interest (AOI). Seventeen saturating pulses were triggered over 8 minutes for a kinetic readout. Due to the ‘Kautsky effect’, only the last three measured values of Y(II) were considered and averaged per leaflet.

### RNA sequencing

Total RNA was extracted from frozen tissue using the PureLink™ RNA Mini kit (ThermoFisher Scientific, Hennigsdorf, Germany) according to manufacturer’s instructions. DNA contamination was removed using the RNAse-free DNAse Set (Qiagen, Hilden, Germany) and additional were removed using the RNeasy^®^ Power Clean^®^ ProCleanup kit (Qiagen). RNA samples of at least 200 ng total RNA and achieving a RIN^e^ greater than 7 on a 4200-tape station (Agilent Technologies) were selected for RNA sequencing. Poly-A capture libraries using the TruSeq Stranded mRNA Library Prep kit were prepared and sequenced by Eurofins Genomics (Konstanz, Germany) on an Illumina NovaSeq 6000 platform in 150 bp paired-end mode for a readout of 30 million read pairs per library.

### RT-qPCR

Four hundred of DNA-free RNA were reverse-transcribed using the qScript™ cDNA SuperMix (Quantabio, Berverly, MA, USA) following manufacturer’s instructions. Quantitative real-time PCR reactions consisted of 12.5 µl PerfeCTa SYBR® Green SuperMix Reagent (Quantabio), 1.25 µl of each primer, 5 µl ddH_2_O, and 5 µl 100-fold diluted cDNA. PCR took place on a qTOWER^3^G cycler (Analytik Jena) with the following parameters: 3 minutes at 95°C and 42 cycles of denaturation for 15 seconds at 95°C and annealing for 30 seconds at 58°C. Melting curve analysis following each PCR run was performed to ensure clean amplification. The efficiency of each primer pair was assessed on 5 cDNA serial dilutions prior to all qPCR events. The following primers were designed based on homology to OrArBC5 sequences from the Parasitic Plant Genome Project (Westwood et al., 2012): HSP17.6-F, GAGGAGAAGAACGACACGTG; HSP17.6-R, TCAGCACTCCGTTCTCCATAC; Ef1α-F, TTGCCGTGAAGGATCTGAAAC; Ef1α-R, CCTTGGCAGGGTCGTCTTTA.

### Reactive oxygen species (ROS) quantification

Determination of reactive oxygen species was carried out as described in (Jambunathan 2010). Briefly, frozen tubercle tissues were ground in liquid nitrogen using sterile mortar and pestle. Ground samples were resuspended in 1 ml Tris-HCl (10 mM) buffer, centrifuged at 12,000 g for 20 minutes at 4°C. The supernatant was transferred into Eppendorf tubes and diluted (1:9) with Tris-HCl (1 mM) buffer. One microliter of 10 mM 2’,7’-Dichlorofluorescein diacetate (DCFH-DA) was added, and 200 µL of the mixture was distributed into 96-well plates. The control consisted of diluted samples without DCFH-DA. All samples were incubated for 10 minutes at room temperature. The ROS were measured using a Infinite M200 TECAN fluorometer (excitation 500 nm, emission 531 nm). Protein content was measured from the same extract using the Pierce™ BCA Protein assay (ThermoFisher Scientific) on a Multiskan Sky High spectrophotometer (Thermo Scientific) at 562 nm absorbance in precision mode.

### Transcriptome analyses

Adaptors and low-quality paired-end sequences were removed from the raw reads using Trimmomatic v0.36 (Bolger et al. 2014). We used Bowtie2 v2.4.1 (Langmead and Salzberg 2012) to map trimmed reads to the OrAeBC5 reference build (http://ppgp.huck.psu.edu) as described earlier (Brun et al. 2023). Read counts per sample were estimated using RSEM v1.3.3 (Li and Dewey 2011).

To identify genes that are differentially expressed during the course of adverse condition treatments and between experimental groups, we performed a comparative time-course differential gene expression analysis using the *maSigPro* R package (Nueda et al. 2014). To this end, read count data was converted to a TMM matrix using the *edgeR* package (Robinson et al. 2010), from which quadratic regression model fitting our four timepoints was computed. Gene-wise regression fits were calculated with an FDR threshold of 0.05 and an F-statistics alpha-level cutoff of 0.05. To identify significantly differentially expressed genes, we subsequently employed a “backward” stepwise regression model with a *P*-value cutoff at 0.05 of the regression coefficients. Significant genes with a minimum R-square of 0.8 of the regression model were extracted by treatment group. Optimal number of clusters was determined using the elbow method by calculating the Within-Cluster-Sum of Squared Errors (WSS). Over-representation analysis of gene ontology (GO) terms was performed based on the previously re-annotated OrAeBC5 build (Brun et al. 2023). Over-representation results and annotation of clustered genes are available as Supplementary Information (Tables S1, 2).

## Results & Discussion

### *Adverse abiotic conditions induce ROS accumulation and* HSP17.6 *expression in* Phelipanche *tubercles*

We examined the physiological responses of the holoparasite *Phelipanche ramosa* to adverse abiotic conditions. To this end, we quantified reactive oxygen species (ROS) contents in parasite tubercles as a marker of stress response due to their documented dual role as signaling molecules in plant growth and as toxic compounds produced through anaerobic metabolic processes during plants abiotic stress responses (Fichman and Mittler 2020). ROS content was expressed as redox-sensitive fluorescing signal (i.e., DCFH-DA fluorescence) relative to total protein abundance in order to account for discrepancies in tubercle weight across samples (Fig. 1). We could not detect significant variations in ROS abundance under control environmental conditions while significant changes in ROS content were observed throughout the kinetic under both adverse heat conditions (*P* < 0.001; Kruskal-Wallis test). Group comparisons through Dunn post hoc test revealed significant five-fold or more increases in ROS content upon heat stress, including after 19.5h of recovery in control conditions. We made similar observations upon high irradiance. Indeed, ROS significantly accumulated by more than 5-fold after 1h of light stress (*P-*adjusted = 0.003) and after recovery (*P-*adjusted = 0.002). More than half the measurements fell above the control median (Fig. 1). The much higher variance observed in light stress conditions may reflect that transient exposure to high irradiance has a milder physiological effect on the parasite, which would be in line with its milder effect on the host photosynthetic status (see below).

**Fig. 1.**
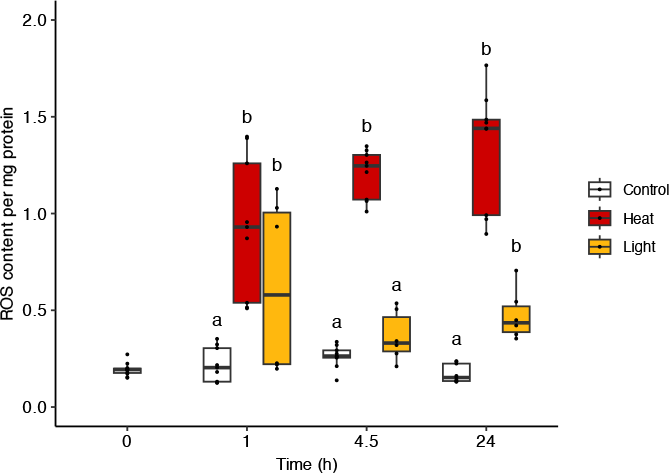
Reactive oxygen species accumulation in *Phelipanche* tubercles upon biotic stressors. Parasite-infested plants were subjected to heat or light stress during 4.5 h and allowed to recover in control conditions for 19.5 h. Reactive oxygen species (ROS) were quantified in *P. ramosa* tubercles and normalized to total protein content. Data are means ± SE (n = 9-12) from two independent experiments. Letters indicate significant differences between environmental conditions (*P-*adjusted < 0.05; Dunn post hoc test followed by Benjamini-Hochberg correction).

Additionally, we investigated the expression of the *Phelipanche* orthologue of the *Arabidopsis HSP17*.*6*, which was previously identified as a marker of heat and high light responses (Sun et al. 2001; Crisp et al. 2017; Huang et al. 2019). Primers were designed based on homology between the *Arabidopsis* sequence (AT5G12030) and the *Phelipanche* reference transcriptome (OrAeBC5 build) from the PPGP database, and expression of the *PrHSP17*.*6* gene was normalized to the expression of the housekeeping gene *PrEF1-*α (Fig. S1). Exposure of the tubercles to heat led to a significant up-regulation by 3.8 ± 0.7 and 2,282 ± 122.5-fold after 1h and 4.5h, respectively. *PrHSP17*.*6* declined to 397.1 ± 90.9-fold after 19.5h of recovery. In contrast, we could not detect any significant variation in gene expression upon light stress (Fig. S1). This corroborates our observations that the impact of transient exposure to elevated temperatures is overall stronger than that of high irradiance.

The parasite is much more affected by transient heat stress compared to high light conditions. RNA sequencing enabled us to cluster genes based on their expression profiles over time and under different treatment conditions. Our analysis found that the quantity of genes differentially expressed in response to heat was clearly greater than in response to light. Finally, the heat wave induced rapid necrosis of the tubercles up to the point where a total loss of viability occurred (Fig. S2).

### Parasite infestation amplifies abiotic stress-induced effects on the host

In parallel, we analyzed the photosynthetic ability of tomato plants under combined biotic and abiotic stress, we used chlorophyll fluorometry to measure the quantum yield of photosystem II (ɸ_PSII_) over time in leaves of tomato plants subjected or not to the holoparasite *Phelipanche* and/or to high irradiance or increased temperatures. Infested and non-infested plants were therefore subjected to abiotic stress for 4.5 h then placed back to control conditions for 19.5 h (Fig. 2). Photosynthesis efficiency was not affected by the presence of the parasite in plants cultivated under control conditions (23°C, 100 µmol m^-2^ s^-1^). Exposure of non-infested plants to elevated temperatures (42°C, 100 µmol m^-2^ s^-1^) led to a two-fold decrease in photosynthesis efficiency after 4.5 h, while recovery in control conditions for 19.5h led to retrieve 89.9 ± 0.9 % of the initial quantum yield. Plants that were infested exhibited a slightly lower photosynthesis efficiency within the first hour of heat stress (*P* = 0.08), and retrieved only 36.1 ± 13.4% of the initial quantum yield after returning to control conditions, indicating a significant failure to recover from heat stress (*P* = 0.004). Quantum yield decreased by about 20 % in non-infested and infested plants after 1 h of light stress, and to a significantly higher extent by 39.2 % in infested plants after 4.5 h (*P =* 0.04). Plants retrieved up to 81.9 ± 0.9 % and 79.6 ± 0.9 % of their initial photosynthesis efficiency after 19.5 h of recovery in control conditions. Altogether these results indicate a negative additive effect of root parasitism and abiotic stresses on host photosynthesis, particularly when plants are subjected to a transient increase in temperature.

**Fig. 2.**
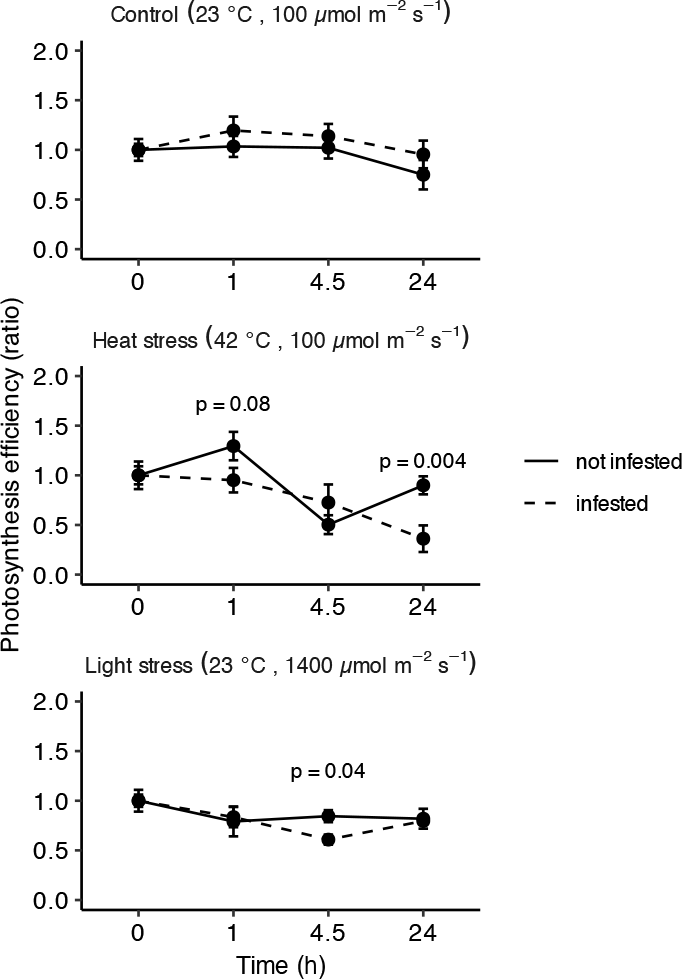
Effect of combined biotic and abiotic stresses on tomato photosynthesis efficiency. Parasite-infested and non-infested tomato plants were subjected to heat or light stress during 4.5 h and allowed to recover in control conditions for 19.5 h. Data are mean ratios ± SE (n = 6-12) from two independent experiments. *P*-values indicate significant differences between infested and non-infested plants at specific timepoints (Kruskal-Wallis test).

### *Transcriptional responses of* Phelipanche *tubercles upon abiotic stresses*

Our physiological analyses indicate that the *Phelipanche* exhibits immediate responses to momentarily adverse temperature and excessive light conditions. This led us to hypothesize that variations in the parasite’s transcriptional landscape are one of the main and immediate modifications. To test this, we collected RNA samples were collected in a time-series experiment to capture transcriptome changes that underlie acclimation and recovery. Trimmed reads were mapped against the broomrape reference transcriptome and the genes were subsequently clustered according to differences in expression profiles over time and between treatment groups.

We determined a total of 1,321 differentially-expressed genes (DEGs), grouped into eight clusters (Fig. 3A) with little functional overlaps between each cluster (Fig. S3). We observed a few variations in gene expression in plants grown under control conditions, mostly occurring in the 4.5-24h window when day/night alternation takes place. Clusters 3, 7 and 8 signify that the broomrape, which relies completely on its host plant for survival, has the genetic makeup to regulate the circadian cycle. Genes associated with the gene ontology (GO) terms GO:0048511 (“rhythmic process”) and GO:0007623 (“circadian clock”) such as CRY1, LNK1, APRR9, and JMJ30 were up-regulated between 4.5 and 24h (Figs. S4A-B, Tables S1-2).

**Fig. 3.**
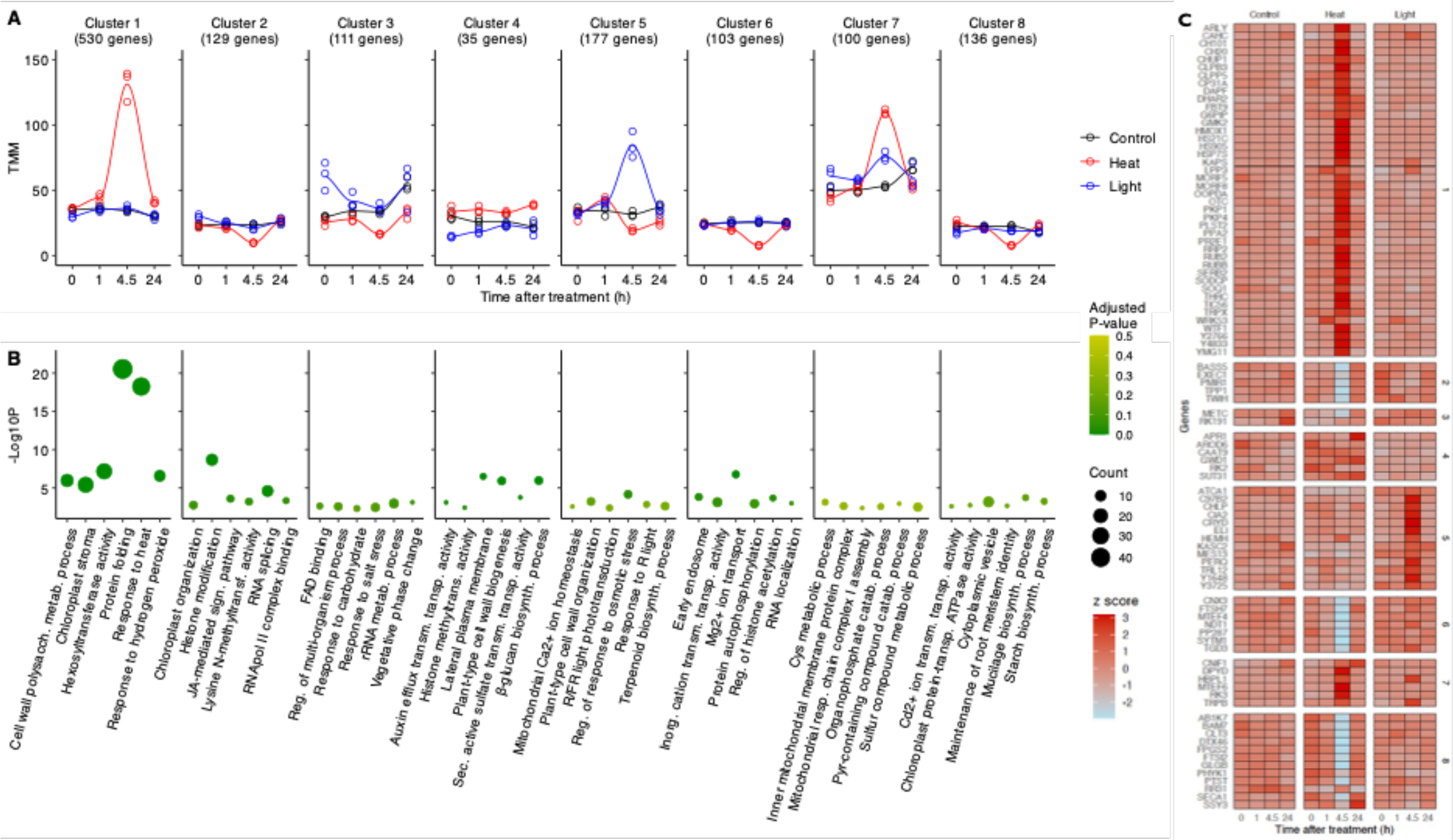
Transcriptional changes upon abiotic stresses in tubercles of *Phelipanche*. **(A)** Gene clusters harboring differential expression patterns of stress response. Parasite-infested plants were subjected to heat or light stress for 4.5h and allowed to recover in control conditions for 19.5 h. Gene expression data were clustered according to their differential accumulation patterns over time and between treatments. The obtained results of significantly differently expressed genes are medians of TMM-normalized counts from three biological replicates. **(B)** Overview of the 6 most significant abbreviated GO terms per cluster, highlighting functional specificity per cluster and treatment. **(C)** A detailed bioinformatic analysis of *Phelipanche*’s plastid-target genes reveals a heat acclimation-specific recruitment of many plastid ribonucleoproteins (cpRNPs), most notably so of CP31A, which is a known integrator of cold acclimation in *Arabidopsis* and whose primary target RNAs no longer exist in the holoparasite’s plastid genome (Wicke et al. 2016).

The overall larger amplitude of expression variation in response to heat represented a significant main difference between abiotic stressors (Fig. 3C). These findings suggest that exposure to high temperatures for a short period of time has a more significant effect than high irradiance. This was particularly evident in clusters 2 and 7, which gathered genes that were predominantly down-regulated and up-regulated in response to 4.5 h of abiotic stress, respectively (Fig. 3). The most enriched biological process associated with cluster 2 was that of “histone modification” (GO:0016570). This included down-regulation of genes encoding histone methyltransferases such as SUVH1 and SUVR2, histone deacetylases from the Sirtuin family such as SIR1 and SNL2, histone acetyltransferases such as IDM1, and histone demethylases such as JMJ25 (Fig. S4C). Noteworthily in cluster 6, additional chromatin remodelers like the histone acetyltransferases ADA2 and the histone deubiquitinase OTU6 were down-regulated upon heat stress only. This overall suggests a certain degree of functional redundancy in epigenetic control in the response to adverse conditions.

Cluster 2 additionally contained genes associated with “jasmonic acid mediated signaling pathways” (GO:0009867), such as RCF3 known as an upstream regulator of heat stress responsive gene expression (Guan et al. 2013), the ubiquitin-specific protease UBP12, and the homeodomain transcription factor OCP3. The latter is known to be involved in drought tolerance (Ding et al. 2013), depending on the perception of jasmonic acid through COI1 (Ramírez et al. 2010) (Fig. S4D). COI1 itself was massively up-regulated upon 4.5h of heat stress together with the ethylene-response factor ERF42. This implies recruitment of the jasmonate-ethylene pathway in response to heat. Another line of evidence is the strong heat-induced down-regulation of the gene encoding NINJA (NOVEL INTERACTOR OF JAZ; Fig. S4D), co-repressing transcription in the absence of jasmonate (Pauwels et al. 2010). On the other hand, biological processes over-represented in cluster 7 were mostly associated with serine and cysteine metabolism. This notably includes up-regulated genes encoding CNIF1 for the maturation of plastidic Fe-S proteins (Léon et al. 2002; Ye et al. 2005), the chloroplast-localized serine acetyltransferase SAT1 for cysteine biosynthesis and known to function in a redox-sensitive module that integrates oxylipin/jasmonate signaling during high light response (Müller et al. 2017), and a cytosolic isoform of cysteine synthase CYSK (Fig. S4E).

Opposite expression profiles between light and heat stresses were mostly found in cluster 5, which included genes highly up-regulated upon 4.5h of light stress while following a milder variation albeit up-down expression pattern upon 1 and 4.5h of heat stress, respectively (Fig. 4). These genes predominantly associated with “response to red light” (GO:0010114) and “red, far-red light phototransduction” (GO:0009585). This included one member of the plastid-localized Early Light-Inducible Proteins (ELI) thought to be involved in photoprotection (Montané and Kloppstech 2000), the bZIP transcription factor HY5, which acts as a positive regulator of phytochrome-mediated photoresponses (Toledo-Ortiz et al. 2014) and anthocyanin accumulation (Shin et al. 2013), the repressor of photomorphogenesis SPA3 thought to be part of the COP1/SPA ubiquitin-protein ligase complex (Laubinger and Hoecker 2003), and the POZ/BTB containing-protein POB1, predicted to be involved in protein ubiquitylation (Figs. S4F-G). Cluster 5 additionally contained genes associated with “terpenoid biosynthetic process” (GO:0016114) and encoding heme-binding proteins such as the geranylgeranyl diphosphate reductase CHLP indirectly involved in tocopherol and chlorophyll synthesis by synthesizing phytol (Tanaka et al. 1999), and the hydroxymethylglutaryl-coA-synthase HMCS (Fig. S4H), which is required for the development of elaioplasts (Ishiguro et al. 2010). Finally, genes belonging to the expansin superfamily were up-regulated in response to either stresses, suggesting a loosening of cell walls upon abiotic stresses. In relation to this, the genes encoding the Leucine-rich repeat receptor-like kinases FEI1 and FEI2, both involved in cell wall biosynthesis, were down-regulated upon heat stress (Fig. S4I).

**Fig. 4.**
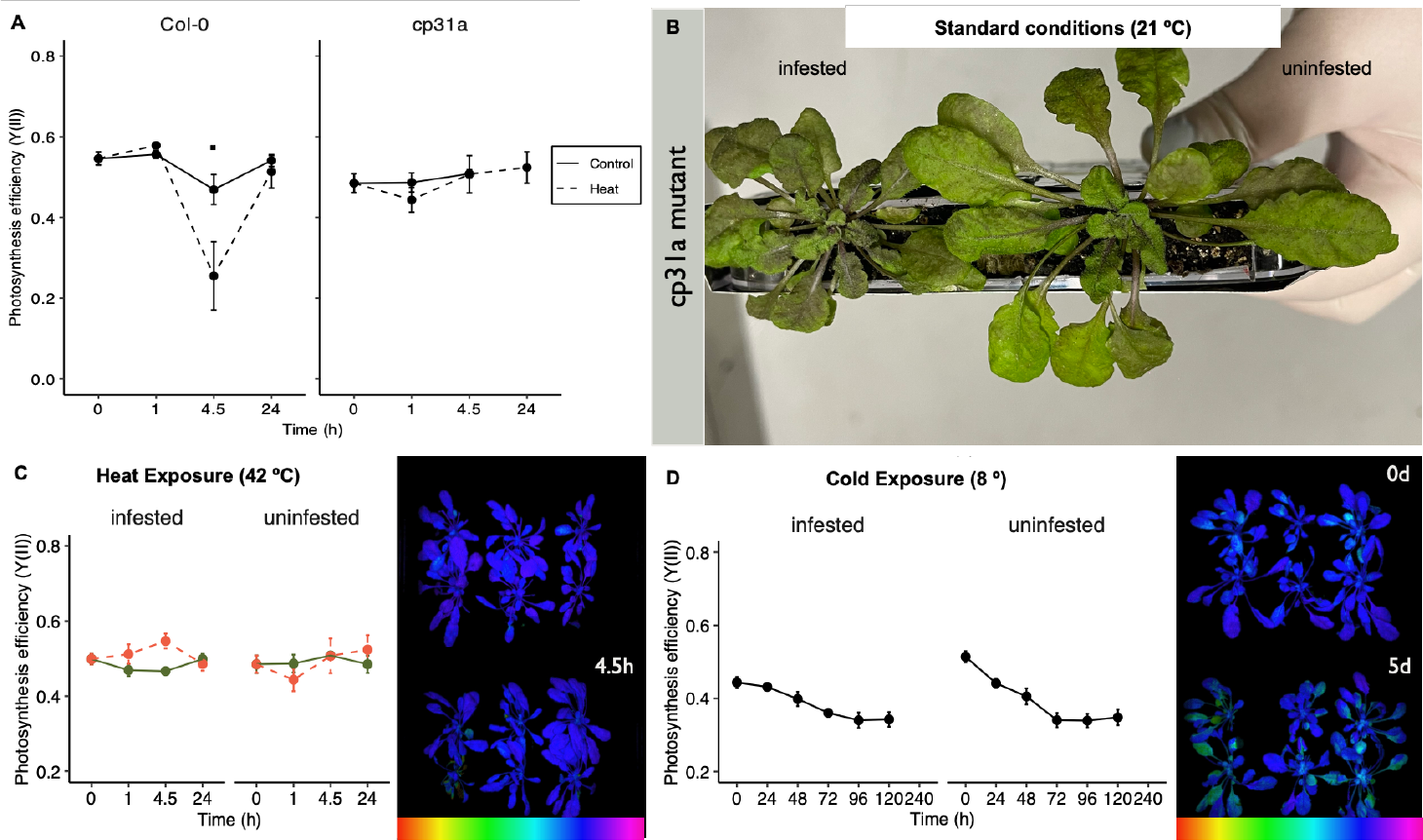
Performance of the *Arabidopsis cp31a* mutant under adverse temperature conditions. **A**. *Arabidopsis* 21-week-old seedlings were subjected to heat stress during 4.5 h and allowed to recover in control conditions for 19.5 h. Data are mean ratios ± SE (n = 3-6) from two independent experiments. Symbols indicate marginal significant differences between treatments at specific timepoints (Kruskal-Wallis test; see text). Parasite infestation in *Arabidopsis* plants **(B)** influences anthocyanin accumulation under standard conditions, **(C)** stabilizes quantum yield around photosystem II (YII) under **(C)** temporary heat conditions and **(D)** over extended cold exposure. Notably, infestation with the parasitic plant reduces acclimation response variability in the cp31A mutant plants. These findings shed light on the complex interplay between biotic (parasite) and abiotic (heat and cold) stress factors and how plants adapt to and synchronize their responses under dual challenges.

Finally, clusters 1 and 6 gathered genes whose expression was exclusively influenced by heat stress however in opposite directions (Fig. 3A). Both clusters shared over-represented biological processes such that of “inorganic cation transmembrane transporter activity” (GO:0022890). Among the up-regulated genes were those encoding Glutamate receptors (GLR) and Na^+^/Ca^2+^ exchangers of the NCL gene family. On the other hand, down-regulated genes were annotated as heavy metal ATPase (HMA) and magnesium transporters (MRS; NIPA) (Fig. S4J). A total of 36 genes up-regulated upon heat stress were annotated as “response to heat” (GO:0009408), more than 80% of which related to families of heat shock proteins (HSP) and heat shock transcription factors (HSF) (Fig. S4K). Of note is the up-regulation of two ClpB/Hsp100 genes known to be essential during chloroplast development, as well as two chloroplast-targeted HSP90 transcripts, known to be crucial for protein import into the stroma (Jiang et al. 2020). Three additional HSP70-encoding genes were also found under the over-represented term “chloroplast stroma” (GO:0009570), along with two plastidial pyruvate kinases, the plastid-localized peroxiredoxin PR2E1, the thylakoid-located thioredoxin-like protein SOQ1, a plastid superoxide dismutase SODM, the plastid phosphoglucose isomerase G6PIP/PGI, and the chloroplast co-chaperonin CH20/CPN21 (Fig. S4L). Finally, we found several genes involved in mitochondria-associated processes, such as the aconitase ACOT9, three chaperonins with homology to the GroES superfamily, the translation factor GUF1, which is involved in mitochondrial protein synthesis (Bauerschmitt et al. 2008), a mitochondrial-processing peptidase MPPB, part of the mitochondrial import complex, and a mitochondrial acyl carrier protein ACPM1, contributing to the respiratory chain.

Altogether, our analysis showed that abiotic stresses such as elevated temperature or light influence the transcriptional landscape of the obligate root parasite *Phelipanche*. Annotation of the differentially-expressed genes indicated that a non-trivial proportion of nuclear-encoded genes influence the function of other major organelles such as mitochondria and chloroplasts, which suggests a major role for anterograde signaling in coping with adverse environmental conditions.

### Plastid-targeted genes are notably differentially upregulated in response to heat

Our data shows that ROS elevation in the parasite (Fig. 1) coincides with the strong up-regulation of many HSP genes (Fig. 3C, Fig. S4K). Interestingly, this includes *HSP21*, which in *Arabidopsis* encodes one of the few proteins that associate with thylakoid membranes upon heat stress (Bernfur et al. 2017). Noteworthily, the holoparasite develops no thylakoid membranes in its achlorophyllous plastids (Laudi and Albertini 1967). Beyond that, the parasite activates genes for plastid-to-nuclear response pathways and the membrane translocation complex (Tic/Toc, CIA2). Apparently, adverse conditions induce the reorganization of the parasite’s plastid structure and relocates them, e.g. by CHUP1 under heat or PMIR1 under light, as our expression data suggests (Fig. 3C, Fig. S5). Several more genes with key function in the plastid are differentially upregulated upon heat exposure, and their identities suggest that plastids of *Phelipanche* retain their primordial roles as environmental sensors. Of the over 460 unigenes associated with plastid-nuclear signaling pathways in our *Phelipanche* reference transcriptome, only an orthologue of HMOX1 is activated after 4.5 h heat treatment (Fig. 3C). This gene is known to encode a plastid heme oxygenase, which is relevant both for phytochrome chromophore biosynthesis and for coupling the expression of selected nuclear genes to the functional state of the plastid. Puzzling are results regarding *Phelipanche*’s heat-specific significant upregulation of chlorophyll- or thylakoid associated pathway genes (e.g. CHLP, SOQ1, RR31; Fig. 3C, Figs. S4K-L) and of the DASH-cryptochrome (CRYD) during recovery (Fig. 3C). In contrast, the differential expression of the uncharacterized proteins Y2766 (copper-iron binding protein At2g37660;), Y4833 (At4g08330), Y3725 (At1g66480), and Y3725 (GTP-binding protein At3g49725) upon heat in the parasite implicates their role in heat response or recovery, respectively, as a first clue to their function. Moreover, heat, rather than high light, strongly induces gene expression to modulate the plastid’s RNA processing machinery in the parasite (Fig. 3).

### Phelipanche *upregulates the RNA-binding protein cp31A in response to heat*

Under heat conditions, the parasite significantly differentially upregulates CP31A (Fig. 3C, alongside other plastid-targeted RBPs, which is most pronounced after 4.5 h of heat exposure. This result is puzzling considering the parasite’s extensive plastid genome reduction. In consequence to *Phelipanche*’s nonphotosynthetic lifestyle, it has lost nearly all plastid-encoded photosynthesis genes including those of the NDH complex, which is the primary target of CP31A (Lenzen et al. 2020). Of CP31A’s previously implicated binding partners (Lenzen et al. 2020), only *matK* as well as *rps2* and *rps8* are encoded in the parasite’s plastome and are expressed under heat (Fig. S5). We may hypothesize that CP31A either targets these genes’ mRNAs, or that the RBP has diversified its specificity for plastid mRNAs or expanded its role beyond cold signal integration in the parasite. Alternatively, it could more generally serve a cryptic function, which has been overshadowed by the canonical key functions of the photosynthetically active chloroplasts in *Arabidopsis*. In addition to cp31A and other RBPs, we find the RNA-binding protein WTF1 to be activated upon heat in *Phelipanche* (Fig. 3C). WTF1 binds to most chloroplast introns, including *trnK, rpl2*, as well as *rps12*, and it is required for splicing of at least *rpl2* (Kroeger et al. 2009). WTF1 might be required to promote splicing the parasite’s three remaining group II introns residing in the genes *trnK, rpl2*, and *rps12* (3’-end); all others have been lost from *Phelipanche* species’ plastomes (Wicke et al. 2016). These findings are puzzling, implying that absence of this plastid-targeted RBPs stabilizes *Arabidopsis*’ heat tolerance. The RBP CP31A is mostly relevant for cold acclimation in *Arabidopsis*. Its upregulation in the holoparasite under heat prompts the question about its general role under temporary heat conditions and which plastid mRNAs CP31A stabilizes in *Phelipanche*.

Chloroplast gene expression is a multifaceted process, commencing with RNA transcription and culminating in a cascade of post-transcriptional events mediated predominantly by RNA-binding proteins (RBPs) (Small et al. 2023). RBPs are encoded in the nucleus and imported into chloroplasts post-translationally. Members of the chloroplast ribonucleoprotein family (cpRNPs) target multiple plastid RNAs and govern chloroplast RNA processing and regulation. The roles of cpRNPs extend from editing and stabilizing RNA to orchestrating chloroplast development in response to environmental fluctuations (Ruwe et al. 2011). To examine the role of *CP31A* in heat response tolerance, we measured photosynthetic ability of *Arabidopsis* Col-0 and *cp31a* mutant seedlings, subjected to heat conditions for 4.5h (Fig. 4). Photosynthesis efficiency significantly decreased by 2.2-fold in the wild-type after 4.5h of heat stress (*P =* 0.06). In contrast, we found no significant alteration of photosynthesis efficiency in the *cp31a* mutant (Fig. 4A). Parasite infestation induces differential anthocyanin accumulation in wildtype *Arabidopsis* plants infested with *Phelipanche* (Fig. 4B). In the presence of parasite infection, both wildtype *Arabidopsis* and all mutant lines exhibit increased stability in the quantum yield of photosystem II (YII) when exposed to heat. The *cp31a* deletion line demonstrates an increase in YII after 4.5 hours of heat exposure (Fig. 4C) and a milder decline in YII under cold conditions over several days (Fig. 4D). This results in a reduced variability in acclimation responses to adverse environmental conditions, suggesting that there is a synchronization of acclimation responses in *Arabidopsis* when faced with dual-challenge conditions.

Together, the newly obtained data suggest several key findings related to acclimation of *Arabidopsis* under dual-challenge conditions. The differential accumulation of anthocyanin in *Arabidopsis* plants indicates that the presence of the parasite triggers a specific response, resulting in a stabilizing effect on the quantum yield of photosystem II (YII) in both wildtype and mutant plants when exposed to adverse heat and cold conditions. The presence of the parasite seems to enhance Arabidopsis’ ability to maintain photosystem II functionality under abiotic stress. The stabilization of photosynthetic efficiency in infected plants leads to a reduction in the variability of acclimation responses to adverse conditions of infected plants, which may be advantageous for their survival. The *cp31a* deletion line exhibits distinct responses under adverse conditions, suggesting that this mutant synchronizes acclimation response, perhaps as part of the plant’s strategy to cope with the combined stressors.

## Conclusions

This study presents the first detailed assessment of acclimation to adverse environmental conditions for a holoparasitic plant. Several studies have shown that parasitic plants of the Orobanchaceae family negatively influence their host’s growth and biomass in correlation with a reduction in host photosynthesis. For example, maximum quantum yield of photosystems was reduced in *Plantago lanceolata* plants infested with the facultative hemiparasite *Rhinantus minor* (Cameron et al. 2008). Similar to this, a steady decrease in transpiration and photosynthesis rates was shown in sorghum varieties that were vulnerable to the obligate hemiparasite *Striga hermonthica* but not in varieties that were partially tolerant to it (Frost et al. 1997). An earlier study also showed that *Striga*-induced reduction in sorghum photosynthesis preferentially occurs before its emergence above ground (Graves et al. 1989). In contrast, our study shows that the obligate holoparasite *Phelipanche ramosa* does not induce any significant reduction in photosynthesis of tomato plants grown under optimal conditions. A sensible interpretation to such contrasted observations may be that the parasite was still several weeks ahead shoot differentiation and aboveground growth. This would be consistent with another study which showed that *Phelipanche* only induced significant and increasingly harmful effects on the initial and maximum fluorescence of chlorophyll in tomatoes after more than a hundred days of interaction, at which time the parasite has long transitioned to reproductive growth (Mauromicale et al. 2008). This leads to assume that the gradual transition of hemiparasites to (semi-)autotrophic growth is accompanied by a relaxed pressure on host photosynthesis, while fully heterotrophic parasites such as weedy broomrapes increasingly suppress host photosynthesis as sink strength amplifies along aboveground, reproductive growth. Significant reduction in host photosynthesis was observed upon momentarily higher temperatures, yet it is likely that the duration of the stress was short enough to allow the tomato plants to recover up to 89% of their initial photosynthesis rates within one day. Heat stress effects were however markedly higher in *Phelipanche*-infested plants both during stress exposure and after recovery. This validates assumptions that combining abiotic and biotic stresses exerts an additively negative influence on host performances. It also implicitly raises numerous concerns as to how C3 plants with poor water use efficiency and high sensitivity to even short exposure to heat and drought stresses will cope with increasingly worrying field infestation rates in the context of global warming.

In contrast, high irradiances did not seem to affect host photosynthesis as significantly. This aligns well with previous reports showing only minor reductions in quantum yield upon brief high light treatment (Anderson et al. 2021) while prolonged or repeated exposures are necessary to influence photosynthesis parameters significantly (Galvez-Valdivieso et al. 2009; Proietti et al. 2023). The same probably holds true for parasite-infested plants, which recovered fully after displaying a significant albeit slight reduction in photosynthesis efficiency upon light stress. Yoshida et al. (2011) demonstrated that rapid acclimation to short-term high light stress in *Arabidopsis* underlies a transient reduction/re-oxidization of the ubiquinone pool concomitant to elevated alternative oxidase levels. This suggests that plants have greater physiological plasticity to light stress than to heat stress in order to maintain their photosynthetic performance, and therefore that there are different degrees of tolerance to parasite infestation depending on both the extent and the nature of unfavorable environmental conditions. The parasite itself also seemed to be much more affected by transient heat stress compared to light stress. Indeed, the increase in ROS content reached a higher level in response to heat and failed to diminish after recovery even after a strong up-regulation of several ROS detoxification marker genes such as peroxidases and superoxide dismutases. In addition, the quantity of genes differentially expressed in response to heat was clearly greater than in response to light. Finally, the heat wave induced rapid necrosis of the tubercles up to the point where a total loss of viability is expected.

In sum, our experiments were the first to highlight that abiotic stress-induced transcriptional changes in an obligate parasitic plant encompass both nuclear and plastid-encoded genes with known implications in plastid-related processes (Fig. 3C, Fig. S5). One such example is the over-representation of magnesium transporters among the heat-repressed genes, which is reminiscent to the central role of Mg^2+^ ions in heat-induced chlorophyll conversion into pheophytin and photosystem degradation (Shimoda et al. 2016). Additional responses to heat stress in autotrophic plants include the maintenance of CO_2_ assimilation capacity as a protective mechanism of Rubisco activity. In tomato, this is facilitated by chloroplast-targeted DnaJ proteins (Kong et al. 2014; Wang et al. 2015), which we also found in multiple copies among the heat-induced genes. The plastid-encoded housekeeping genes *clpP, accD*, and *rps14* were reported to be down-regulated upon heat in *Arabidopsis* (Danilova et al. 2018), which is consistent with the expression patterns of the corresponding orthologs in *Phelipanche*. Therewith, the present study highlights that abiotic stress-induced transcriptional changes in obligate parasitic plants encompass both nuclear and plastid-encoded genes with known implications in plastid-related processes.

## Acknowledgments

The authors thank Christian Schmitz-Linneweber, Kerstin Kaufmann, and Bernhard Grimm and their respective research teams at Humboldt-Universität zu Berlin for assistance regarding technical and other lab equipment. Thanks are also due to Dr. Jean-Bernard Pouvreau (Nantes University, France) for sharing seeds of *Phelipanche ramosa*. This study received financial support by the Deutsche Forschungsgemeinschaft (DFG, WI4507/3-1 and CRC TRR175-A08 to S.W.), which is gratefully acknowledged.

## Data Availability

The data underlying this article are available in Mendeley Data, doi: 10.17632/5w8z7wgnc3, and the NCBI Short Read Archive, BioProject ID: PRJNA1000199.

## Author contributions

Conceptualization: S.W.; Research coordination: G.B. and S.W.; Experiments: O.D., G.B., C.G.; Data analysis: O.D., G.B., C.G., S.W.; Writing & Reviewing: O.D., G.B., C.G., S.W.

## References

Anderson CM, Mattoon EM, Zhang N, Becker E, McHargue W, Yang J, Patel D, Dautermann O, McAdam SAM, Tarin T, Pathak S, Avenson TJ, Berry J, Braud M, Niyogi KK, Wilson M, Nusinow DA, Vargas R, Czymmek KJ, Eveland AL, Zhang R (2021) High light and temperature reduce photosynthetic efficiency through different mechanisms in the C4 model Setaria viridis. Commun Biol 4: 1092

Bauerschmitt H, Funes S, Herrmann JM (2008) The membrane-bound GTPase Guf1 promotes mitochondrial protein synthesis under suboptimal conditions. J Biol Chem 283: 17139–17146

Beltrán J, Wamboldt Y, Sanchez R, LaBrant EW, Kundariya H, Virdi KS, Elowsky C, Mackenzie SA (2018) Specialized plastids trigger tissue-specific signaling for systemic stress response in plants. Plant Physiol 178: 672–683

Bernfur K, Rutsdottir G, Emanuelsson C (2017) The chloroplast-localized small heat shock protein Hsp21 associates with the thylakoid membranes in heat-stressed plants. Protein Sci 26: 1773–1784

Bolger AM, Lohse M, Usadel B (2014) Trimmomatic: a flexible trimmer for Illumina sequence data. Bioinformatics 30: 2114–2120

Brun G, Leman JKH, Wicke S (2023) Comparative gene expression analysis of differentiated terminal and lateral haustoria of the obligate root parasitic plant Phelipanche ramosa (Orobanchaceae). Plants People Planet doi.org/10.1002/ppp3.10464

Cameron DD, Geniez J-M, Seel WE, Irving LJ (2008) Suppression of host photosynthesis by the parasitic plant Rhinanthus minor. Ann Bot 101: 573–578

Chen X, Fang D, Wu C, Liu B, Liu Y, Sahu SK, Song B, Yang S, Yang T, Wei J, Wang X, Zhang W, Xu Q, Wang H, Yuan L, Liao X, Chen L, Chen Z, Yuan F, Chang Y, Lu L, Yang H, Wang J, Xu X, Liu X, Wicke S, Liu H (2020) Comparative plastome analysis of root- and stem-feeding parasites of Santalales untangle the footprints of feeding mode and lifestyle transitions. Genome Biol Evol 12: 3663–3676

Choudhury FK, Rivero RM, Blumwald E, Mittler R (2017) Reactive oxygen species, abiotic stress and stress combination. Plant J 90: 856–867

Crisp PA, Ganguly DR, Smith AB, Murray KD, Estavillo GM, Searle I, Ford E, Bogdanović O, Lister R, Borevitz JO, Eichten SR, Pogson BJ (2017) Rapid recovery gene downregulation during excess-light stress and recovery in Arabidopsis. Plant Cell 29: 1836–1863

Crosatti C, Rizza F, Badeck FW, Mazzucotelli E, Cattivelli L (2013) Harden the chloroplast to protect the plant. Physiol Plant 147: 55–63

Cui S, Kubota T, Nishiyama T, Ishida JK, Shigenobu S, Shibata TF, Toyoda A, Hasebe M, Shirasu K, Yoshida S (2020) Ethylene signaling mediates host invasion by parasitic plants. Sci Adv 6: eabc2385

Danilova MN, Kudryakova NV, Andreeva AA, Doroshenko AS, Pojidaeva ES, Kusnetsov VV (2018) Differential impact of heat stress on the expression of chloroplast-encoded genes. Plant Physiol Biochem 129: 90–100

Ding Y, Liu N, Virlouvet L, Riethoven J-J, Fromm M, Avramova Z (2013) Four distinct types of dehydration stress memory genes in Arabidopsis thaliana. BMC Plant Biol 13: 229

Dopp IJ, Yang X, Mackenzie SA (2021) A new take on organelle-mediated stress sensing in plants. New Phytol 230: 2148–2153

Fichman Y, Mittler R (2020) Rapid systemic signaling during abiotic and biotic stresses: is the ROS wave master of all trades? Plant J 102: 887–896

Frost DL, Gurney AL, Press MC, Scholes JD (1997) Striga hermonthica reduces photosynthesis in sorghum: the importance of stomatal limitations and a potential role for ABA? Plant Cell & Environment 20: 483–492

Galvez-Valdivieso G, Fryer MJ, Lawson T, Slattery K, Truman W, Smirnoff N, Asami T, Davies WJ, Jones AM, Baker NR, Mullineaux PM (2009) The high light response in Arabidopsis involves ABA signaling between vascular and bundle sheath cells. Plant Cell 21: 2143–2162

Graves JD, Press MC, Stewart GR (1989) A carbon balance model of the sorghum-Striga hermonthica host-parasite association. Plant Cell & Environment 12: 101–107

Guan Q, Wen C, Zeng H, Zhu J (2013) A KH domain-containing putative RNA-binding protein is critical for heat stress-responsive gene regulation and thermotolerance in Arabidopsis. Mol Plant 6: 386–395

Huang J, Zhao X, Chory J (2019) The Arabidopsis transcriptome responds specifically and dynamically to high light stress. Cell Rep 29: 4186-4199.e3

Ichihashi Y, Mutuku JM, Yoshida S, Shirasu K (2015) Transcriptomics exposes the uniqueness of parasitic plants. Brief Funct Genomics 14: 275–282

Ishiguro S, Nishimori Y, Yamada M, Saito H, Suzuki T, Nakagawa T, Miyake H, Okada K, Nakamura K (2010) The Arabidopsis FLAKY POLLEN1 gene encodes a 3-hydroxy-3-methylglutaryl-coenzyme A synthase required for development of tapetum-specific organelles and fertility of pollen grains. Plant Cell Physiol 51: 896–911

Jambunathan N (2010) Determination and detection of reactive oxygen species (ROS), lipid peroxidation, and electrolyte leakage in plants. Methods Mol Biol 639: 292–298

Jiang T, Mu B, Zhao R (2020) Plastid chaperone HSP90C guides precursor proteins to the SEC translocase for thylakoid transport. J Exp Bot 71: 7073–7087

Jones JDG, Dangl JL (2006) The plant immune system. Nature 444: 323–329

Kleine T, Nägele T, Neuhaus HE, Schmitz-Linneweber C, Fernie AR, Geigenberger P, Grimm B, Kaufmann K, Klipp E, Meurer J, Möhlmann T, Mühlhaus T, Naranjo B, Nickelsen J, Richter A, Ruwe H, Schroda M, Schwenkert S, Trentmann O, Willmund F, Zoschke R, Leister D (2021) Acclimation in plants - the Green Hub consortium. Plant J 106: 23–40

Kong F, Deng Y, Wang G, Wang J, Liang X, Meng Q (2014) LeCDJ1, a chloroplast DnaJ protein, facilitates heat tolerance in transgenic tomatoes. J Integr Plant Biol 56: 63–74

Kroeger TS, Watkins KP, Friso G, van Wijk KJ, Barkan A (2009) A plant-specific RNA-binding domain revealed through analysis of chloroplast group II intron splicing. Proc Natl Acad Sci U S A 106: 4537–4542

Langmead B, Salzberg SL (2012) Fast gapped-read alignment with Bowtie 2. Nat Methods 9: 357–359

Laubinger S, Hoecker U (2003) The SPA1-like proteins SPA3 and SPA4 repress photomorphogenesis in the light. Plant J 35: 373–385

Laudi G, Albertini A (1967) Infrastructural researches on the plastids of parasitic plants. III. Orobanche ramosa. Caryologia 20: 207–216

Lenzen B, Rühle T, Lehniger M-K, Okuzaki A, Labs M, Muino JM, Ohler U, Leister D, Schmitz-Linneweber C (2020) The chloroplast RNA binding protein CP31A has a preference for mRNAs encoding the subunits of the chloroplast NAD(P)H dehydrogenase complex and Is required for their accumulation. Int J Mol Sci 21: 5633

Léon S, Touraine B, Briat J-F, Lobréaux S (2002) The AtNFS2 gene from Arabidopsis thaliana encodes a NifS-like plastidial cysteine desulphurase. Biochem J 366: 557–564

Li B, Dewey CN (2011) RSEM: accurate transcript quantification from RNA-Seq data with or without a reference genome. BMC Bioinformatics 12: 323

Lyko P, Wicke S (2021) Genomic reconfiguration in parasitic plants involves considerable gene losses alongside global genome size inflation and gene births. Plant Physiol 186: 1412–1423

Mauromicale G, Monaco AL, Longo AMG (2008) Effect of Branched Broomrape (Orobanche ramosa) infection on the growth and photosynthesis of tomato. Weed sci 56: 574–581

Montané MH, Kloppstech K (2000) The family of light-harvesting-related proteins (LHCs, ELIPs, HLIPs): was the harvesting of light their primary function? Gene 258: 1–8

Müller SM, Wang S, Telman W, Liebthal M, Schnitzer H, Viehhauser A, Sticht C, Delatorre C, Wirtz M, Hell R, Dietz K-J (2017) The redox-sensitive module of cyclophilin 20-3, 2-cysteine peroxiredoxin and cysteine synthase integrates sulfur metabolism and oxylipin signaling in the high light acclimation response. Plant J 91: 995–1014

Nueda MJ, Tarazona S, Conesa A (2014) Next maSigPro: updating maSigPro bioconductor package for RNA-seq time series. Bioinformatics 30: 2598–2602

Pandey P, Ramegowda V, Senthil-Kumar M (2015) Shared and unique responses of plants to multiple individual stresses and stress combinations: physiological and molecular mechanisms. Front Plant Sci 6: 723

Pauwels L, Barbero GF, Geerinck J, Tilleman S, Grunewald W, Pérez AC, Chico JM, Bossche RV, Sewell J, Gil E, García-Casado G, Witters E, Inzé D, Long JA, De Jaeger G, Solano R, Goossens A (2010) NINJA connects the co-repressor TOPLESS to jasmonate signalling. Nature 464: 788–791

Proietti S, Paradiso R, Moscatello S, Saccardo F, Battistelli A (2023) Light intensity affects the assimilation rate and carbohydrates partitioning in spinach grown in a controlled environment. Plants (Basel) 12: 804

Ramírez V, Van der Ent S, García-Andrade J, Coego A, Pieterse CMJ, Vera P (2010) OCP3 is an important modulator of NPR1-mediated jasmonic acid-dependent induced defenses in Arabidopsis. BMC Plant Biol 10: 199

Robinson MD, McCarthy DJ, Smyth GK (2010) edgeR: a Bioconductor package for differential expression analysis of digital gene expression data. Bioinformatics 26: 139–140

Ruwe H, Kupsch C, Teubner M, Schmitz-Linneweber C (2011) The RNA-recognition motif in chloroplasts. J Plant Physiol 168: 1361–1371

Serrano I, Audran C, Rivas S (2016) Chloroplasts at work during plant innate immunity. J Exp Bot 67: 3845–3854

Shimoda Y, Ito H, Tanaka A (2016) Arabidopsis STAY-GREEN, Mendel’s green cotyledon gene, encodes magnesium-dechelatase. Plant Cell 28: 2147–2160

Shin DH, Choi M, Kim K, Bang G, Cho M, Choi S-B, Choi G, Park Y-I (2013) HY5 regulates anthocyanin biosynthesis by inducing the transcriptional activation of the MYB75/PAP1 transcription factor in Arabidopsis. FEBS Lett 587: 1543–1547

Small I, Melonek J, Bohne A-V, Nickelsen J, Schmitz-Linneweber C (2023) Plant organellar RNA maturation. Plant Cell 35: 1727–1751

Sun G, Xu Y, Liu H, Sun T, Zhang J, Hettenhausen C, Shen G, Qi J, Qin Y, Li J, Wang L, Chang W, Guo Z, Baldwin IT, Wu J (2018) Large-scale gene losses underlie the genome evolution of parasitic plant Cuscuta australis. Nat Commun 9: 2683

Sun W, Bernard C, van de Cotte B, Van Montagu M, Verbruggen N (2001) At-HSP17.6A, encoding a small heat-shock protein in Arabidopsis, can enhance osmotolerance upon overexpression. Plant J 27: 407–415

Tanaka R, Oster U, Kruse E, Rüdiger W, Grimm B (1999) Reduced activity of geranylgeranyl reductase leads to loss of chlorophyll and tocopherol and to partially geranylgeranylated chlorophyll in transgenic tobacco plants expressing antisense RNA for geranylgeranyl reductase1. Plant Physiol 120: 695–704

Tillich M, Hardel SL, Kupsch C, Armbruster U, Delannoy E, Gualberto JM, Lehwark P, Leister D, Small ID, Schmitz-Linneweber C (2009) Chloroplast ribonucleoprotein CP31A is required for editing and stability of specific chloroplast mRNAs. Proc Natl Acad Sci U S A 106: 6002–6007

Toledo-Ortiz G, Johansson H, Lee KP, Bou-Torrent J, Stewart K, Steel G, Rodríguez-Concepción M, Halliday KJ (2014) The HY5-PIF regulatory module coordinates light and temperature control of photosynthetic gene transcription. PLoS Genet 10: e1004416

Vogel A, Schwacke R, Denton AK, Usadel B, Hollmann J, Fischer K, Bolger A, Schmidt MH-W, Bolger ME, Gundlach H, Mayer KFX, Weiss-Schneeweiss H, Temsch EM, Krause K (2018) Footprints of parasitism in the genome of the parasitic flowering plant Cuscuta campestris. Nat Commun 9: 2515

Wang G, Kong F, Zhang S, Meng X, Wang Y, Meng Q (2015) A tomato chloroplast-targeted DnaJ protein protects Rubisco activity under heat stress. J Exp Bot 66: 3027–3040

Wicke S, Müller KF, dePamphilis CW, Quandt D, Bellot S, Schneeweiss GM (2016) Mechanistic model of evolutionary rate variation en route to a nonphotosynthetic lifestyle in plants. Proc Natl Acad Sci USA 113: 9045–9050

Wicke S, Müller KF, dePamphilis CW, Quandt D, Wickett NJ, Zhang Y, Renner SS, Schneeweiss GM (2013) Mechanisms of functional and physical genome reduction in photosynthetic and non-photosynthetic parasitic plants of the broomrape family. Plant Cell 25: 3711–3725

Wicke S, Naumann J (2018) Molecular evolution of plastid genomes in parasitic flowering plants. In: Chaw S-M, Jansen RK (eds) Plastid genome evolution, 1st Ed. Elsevier, pp. 315–347

Xu Y, Zhang J, Ma C, Lei Y, Shen G, Jin J, Eaton DAR, Wu J (2022) Comparative genomics of orobanchaceous species with different parasitic lifestyles reveals the origin and stepwise evolution of plant parasitism. Mol Plant 15: 1384–1399

Yamori W, Sakata N, Suzuki Y, Shikanai T, Makino A (2011) Cyclic electron flow around photosystem I via chloroplast NAD(P)H dehydrogenase (NDH) complex performs a significant physiological role during photosynthesis and plant growth at low temperature in rice. Plant J Epup ahead of print:

Yang Z, Wafula EK, Honaas LA, Zhang H, Das M, Fernandez-Aparicio M, Huang K, Bandaranayake PCG, Wu B, Der JP, Clarke CR, Ralph PE, Landherr L, Altman NS, Timko MP, Yoder JI, Westwood JH, dePamphilis CW (2015) Comparative transcriptome analyses reveal core parasitism genes and suggest gene duplication and repurposing as sources of structural novelty. Mol Biol Evol 32: 767–790

Ye H, Garifullina GF, Abdel-Ghany SE, Zhang L, Pilon-Smits EAH, Pilon M (2005) The chloroplast NifS-like protein of Arabidopsis thaliana is required for iron-sulfur cluster formation in ferredoxin. Planta 220: 602–608

Yoshida K, Watanabe CK, Hachiya T, Tholen D, Shibata M, Terashima I, Noguchi K (2011) Distinct responses of the mitochondrial respiratory chain to long- and short-term high-light environments in Arabidopsis thaliana. Plant Cell & Environment 34: 618–628

Yoshida S, Kim S, Wafula EK, Tanskanen J, Kim Y-M, Honaas L, Yang Z, Spallek T, Conn CE, Ichihashi Y, Cheong K, Cui S, Der JP, Gundlach H, Jiao Y, Hori C, Ishida JK, Kasahara H, Kiba T, Kim M-S, Koo N, Laohavisit A, Lee Y-H, Lumba S, McCourt P, Mortimer JC, Mutuku JM, Nomura T, Sasaki-Sekimoto Y, Seto Y, Wang Y, Wakatake T, Sakakibara H, Demura T, Yamaguchi S, Yoneyama K, Manabe R-I, Nelson DC, Schulman AH, Timko MP, dePamphilis CW, Choi D, Shirasu K (2019) Genome sequence of Striga asiatica provides insight into the evolution of plant parasitism. Curr Biol 29: 3041–3052

