## Supplementary material for "Acclimation kinetics of the holoparasitic weed *Phelipanche ramosa* (Orobanchaceae) during excessive light and heat conditions": Fig. S1

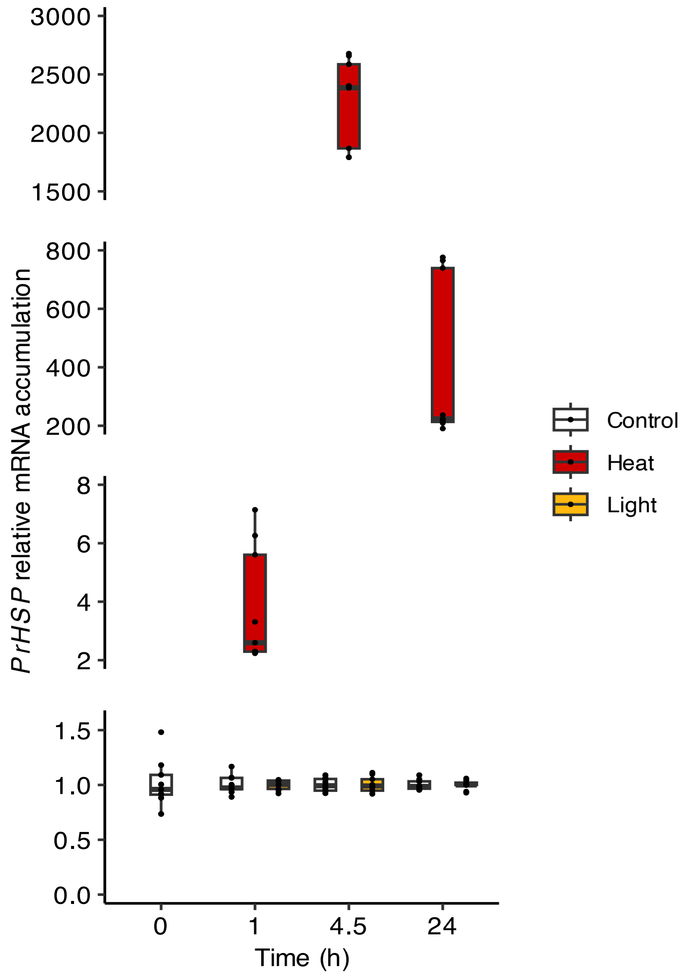


**Fig. S1. *HSP17.6* gene expression in *P. ramosa* tubercles upon adverse abiotic conditons.** Parasite-tomato pathosystems were subjected to heat or light stress during 4.5 h and allowed to recover in control conditions for 19.5 h. Expression of the *PrHSP17.6* gene was quantified by RT-qPCR in *Phelipanche* tubercles and normalized to the expression of the *PrEF1*-α housekeeping gene. Data are means ± standard error (n = 9-18) from two independent experiments.

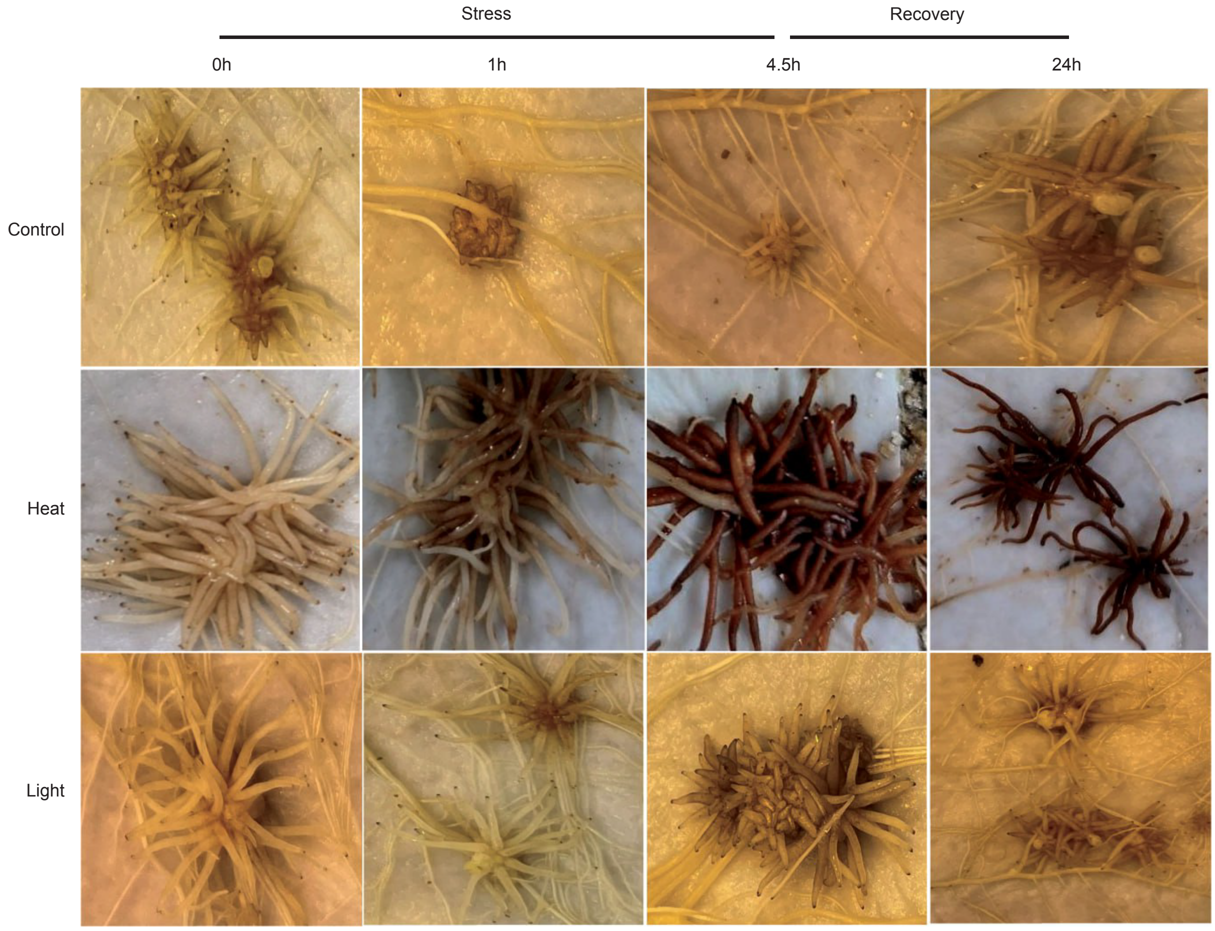


**Fig. S2. Tubercle development upon exposure to heat and excessive light.** Representative pictures of spider tubercles grown on tomato roots for approximately 5 weeks before exposure to 4.5 hours of heat or light stress and recovery for 19.5h.

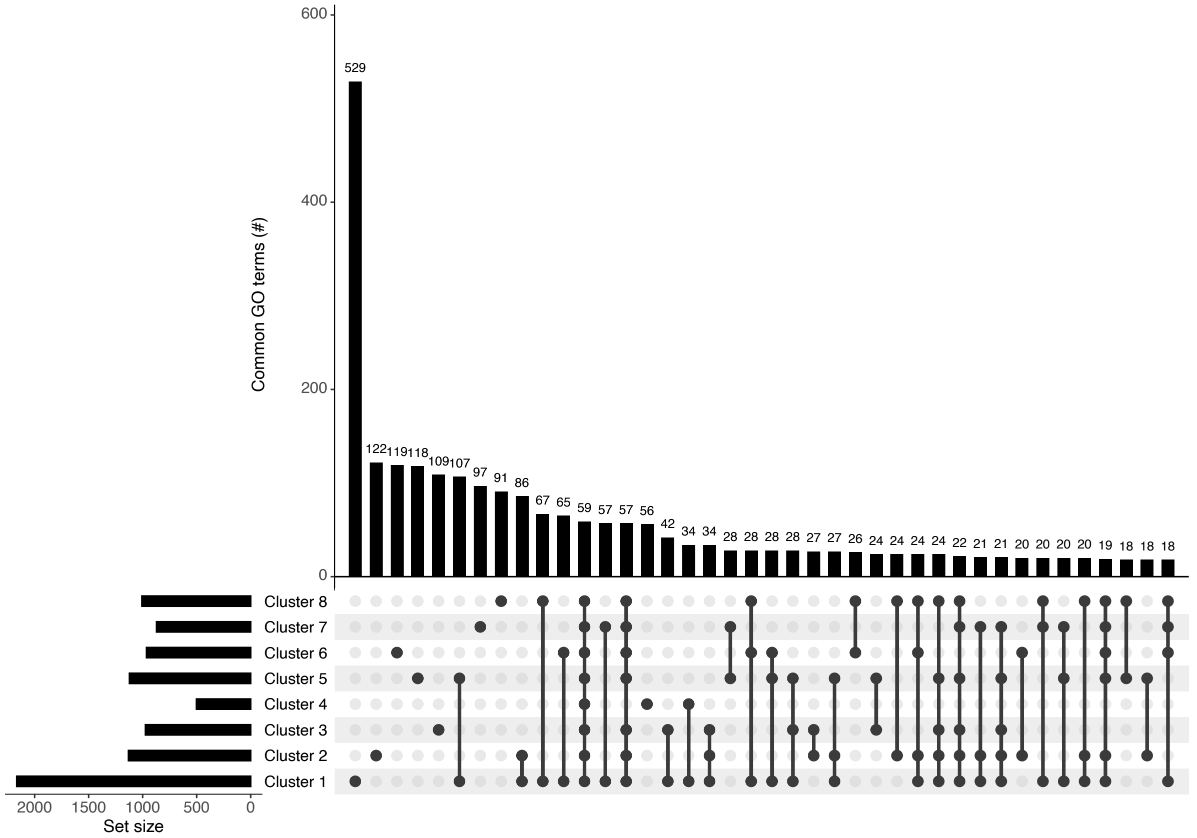


**Fig. S3. Functional differences across gene clusters.** Upset plot summarizing the intersecting terms of all GO categories between clusters (see Fig. 3).

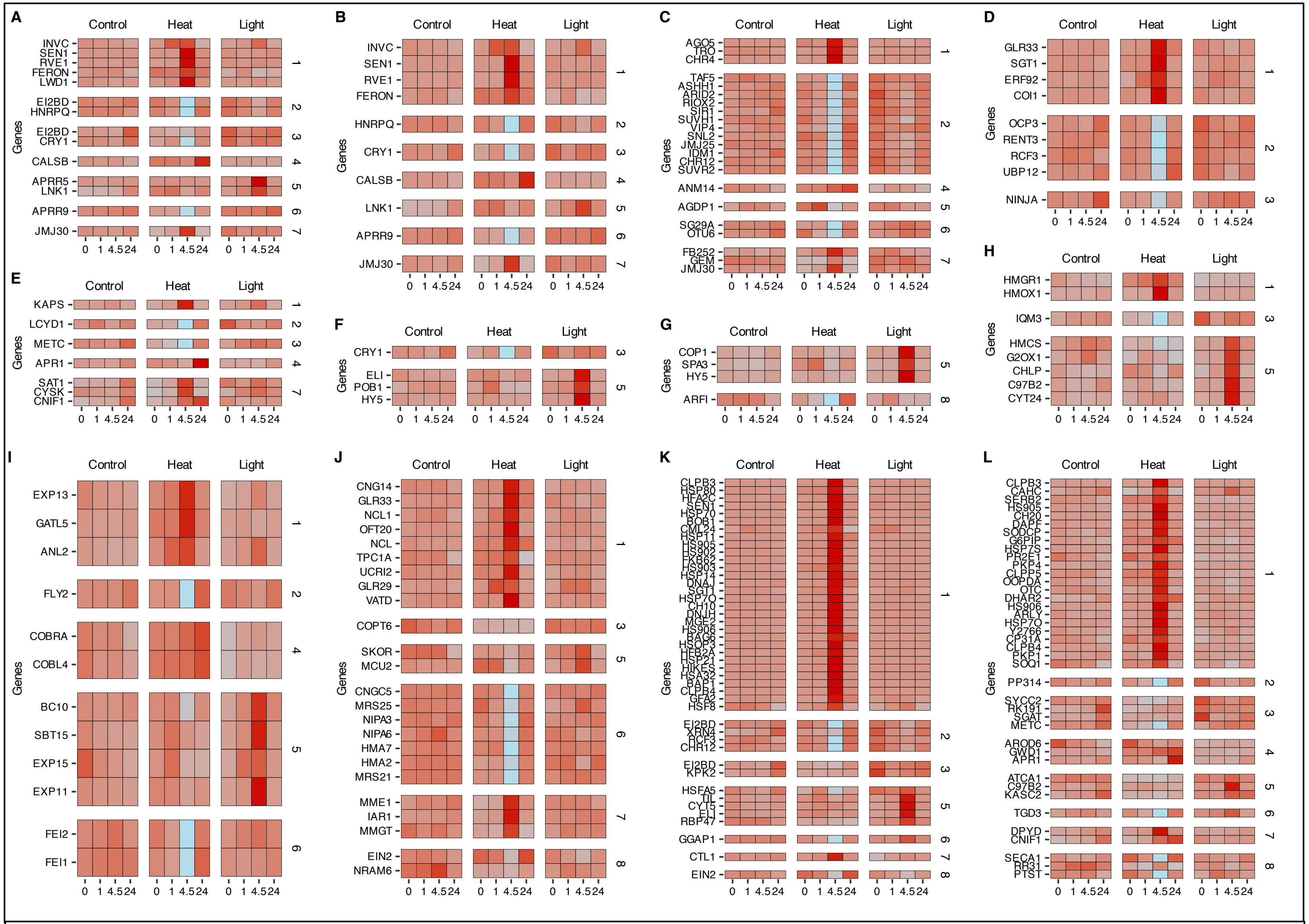


**Fig. S4. Gene expression of over-represented GO terms.** Data are means of Z-scored values for the genes annotated to “rhythmic process” (A), “circadian clock” (B), “histone modification” (C), “JA-mediated signaling pathway” (D), “cysteine metabolic process” (E), “response to ref light” (F), “R/FR light photransduction” (G), “terpenoid biosynthetic process” (H), “inorganic cation transmembrane transporter activity” (I), “response to heat” (J), and “chloroplast stroma” (K).

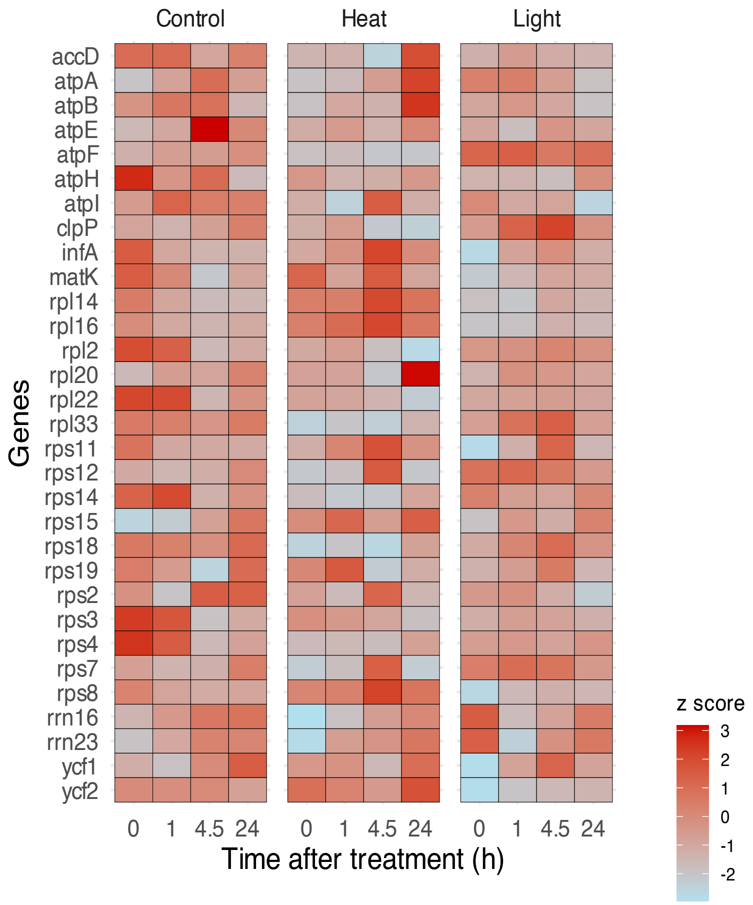


**Fig. S5. Plastid gene expression in *Phelipanche ramosa* tubercles upon exposure to heat and excessive light.** Trimmed reads were mapped onto the remaining plastid genes of *P. ramosa*. Data are means of Z-scored TMM-normalized values.
